# Development of a Novel CD4^+^ TCR Transgenic Line that Reveals a Dominant Role for CD8^+^ DC and CD40-Signaling in the Generation of Helper and CTL Responses to Blood Stage Malaria

**DOI:** 10.1101/113837

**Authors:** Daniel Fernandez-Ruiz, Lei Shong Lau, Nazanin Ghazanfari, Claerwen M Jones, Wei Yi Ng, Gayle M Davey, Dorothee Berthold, Lauren Holz, Yu Kato, Ganchimeg Bayarsaikhan, Sanne H. Hendriks, Kylie R James, Anton Cozijnsen, Vanessa Mollard, Tania F de Koning-Ward, Paul R Gilson, Tsuneyasu Kaisho, Ashraful Haque, Brendan S Crabb, Francis R Carbone, Geoffrey I. McFadden, William R Heath

**Affiliations:** Department of Microbiology and Immunology, Peter Doherty Institute for Infection and Immunity, University of Melbourne, Victoria 3000, Australia; The ARC Centre of Excellence in Advanced Molecular Imaging, University of Melbourne, Parkville, Victoria 3010, Australia.; Division of Immunology, Department of Molecular Microbiology and Immunology, Graduate School of Biomedical Sciences, Nagasaki University, Sakamoto, Nagasaki 852-8523, Japan; Malaria Immunology Laboratory, QIMR Berghofer Medical Research Institute, Herston, Queensland 4006, Australia; The School of BioSciences, University of Melbourne, Parkville, Victoria 3010, Australia; School of Medicine, Deakin University, Waurn Ponds, Victoria 3216, Australia; Macfarlane Burnet Institute for Medical Research & Public Health, Victoria 3004, Australia; Department of Immunology, Institute of Advanced Medicine, Wakayama Medical University, Wakayama, Wakayama 641-8509, Japan

## Abstract

We describe an MHC II (IA^b^)-restricted T cell receptor (TCR) transgenic mouse line that produces CD4^+^ T cells specific for *Plasmodium* species. This line, termed PbT-II, was derived from a CD4^+^ T cell hybridoma generated to blood-stage *Plasmodium berghei* ANKA (PbA). PbT-II cells responded to all *Plasmodium* species and stages tested so far, including rodent (PbA, *P. berghei* NK65, *P. chabaudi* AS and *P. yoelii* 17XNL) and human (*P*. *falciparum)* blood-stage parasites as well as irradiated PbA sporozoites. PbT-II cells can provide help for generation of antibody to *P. chabaudi* infection and can control this otherwise lethal infection in CD40L-deficient mice. PbT-II cells can also provide help for development of CD8^+^ T cell-mediated experimental cerebral malaria (ECM) during PbA infection. Using PbT-II CD4+ T cells and the previously described PbT-I CD8^+^ T cells, we determined the dendritic cell (DC) subsets responsible for immunity to PbA blood-stage infection. CD8^+^ DC (a subset of XCR1^+^ DC) were the major antigen presenting cell (APC) responsible for activation of both T cell subsets, though other DC also contributed to CD4^+^ T cell responses. Depletion of CD8^+^ DC at the beginning of infection prevented ECM development and impaired both Th1 and Tfh responses; in contrast, late depletion did not affect ECM. This study describes a novel and versatile tool for examining CD4^+^ T cell immunity during malaria and provides evidence that CD4^+^ T cell help, acting via CD40L signalling, can promote immunity or pathology to blood stage malaria largely through antigen presentation by CD8^+^ DC.

## Introduction

Despite intervention strategies, malaria killed almost half a million people in 2015 (1). Murine models for malaria present similarities with human infections and allow for the detailed study of immunological processes of potential relevance to human disease (2-8). T cell receptor (TCR) transgenic murine lines specific for pathogen-derived antigens are powerful tools for studying the mechanisms involved in the development of immune responses during infection. Their ease of use and potential for manipulation offers a much broader range of opportunities for the study of T cell responses than are feasible using the endogenous T cell repertoire.

The lack of TCR transgenic mouse lines specific for *Plasmodium* antigens led to the generation of transgenic malaria parasites expressing model antigens, such as PbTG and OVA-PbA (2, 4, 9, 10), for which widely used murine T cell lines such as OT-I and OT-II could be used to monitor specific T cell responses. While the use of these parasites in conjunction with model T cell lines has aided the study of anti-malarial CD4^+^ and CD8^+^ T cell responses (6, 11-15), wild-type parasites and transgenic T cells capable of recognizing authentic parasite-derived antigens are preferred as they more closely resemble endogenous responses to natural infections. With this in mind, we recently generated a murine T cell receptor (TCR) transgenic line of *P. berghei* ANKA (PbA)-specific CD8^+^ T cells, termed PbT-I (8, 16). Here, we introduce a new MHC II-restricted (IA^b^) TCR murine line, termed PbT-II, that responds to a parasite antigen expressed across multiple rodent and human *Plasmodium* species, making it a general tool for studying malaria immunity in C57BL/6 (B6) mice. PbT-II TCR transgenic mice add to the existing I-E^d^-restricted B5 TCR transgenic mice (2, 4, 17) to extend the set of available tools for the analysis of CD4^+^ T cell responses to parasites during *Plasmodium* infection of B6 mice.

CD4^+^ T cells orchestrate both humoral and cellular adaptive immune responses against pathogens. Cross-talk between CD4^+^ T cells and naïve B cells resulting in Ig class switching is essential for the clearance of certain pathogens such as *P. chabaudi* AS. Thus, mice lacking CD4^+^ T cells or B cells are unable to control parasitaemia in this model (17). Another important role for CD4^+^ T cells is the provision of help resulting in the licensing of dendritic cells (DC) for the effective priming of CD8^+^ T cells. However, while CD4^+^ T cell help is essential for primary responses to certain pathogens, such as herpes simplex virus (HSV) (11, 18), it is dispensable during infection with influenza A virus, lymphocytic choriomeningitis virus or *Listeria monocytogenes* (14, 19-21). It is understood that in the latter cases, sufficient engagement of receptors for pathogen associated molecular patterns (PAMP) on DC by material derived from the infectious agent (6, 22), or cytokines secreted by innate cells upon recognition of the pathogen (23, 24), bypass the need for CD4+ T cell help. In the case of PbA infection, the helper dependence of CD8^+^ T cell responses has not been directly addressed. PbA infection of B6 mice leads to the development of experimental cerebral malaria (ECM), a pathology mediated by CD8^+^ T cells that is used as a model for human cerebral malaria (25). Therefore, dissection of the mechanisms that lead to CD8^+^ T cell activation in this model is of importance to better understand this pathology. ECM was abolished when CD4^+^ T cells were depleted during the early stages of the infection in a number of studies (26-30), suggesting that CD8^+^ T cell priming relies on CD4^+^ T cell help. Indeed, depletion of CD4^+^ T cells during infection with the transgenic malaria parasite PbTG resulted in diminished OT-I cell proliferation (31). However, transfer of OT-I cells into PbTG-infected RAG mice, lacking T cells, resulted in the development of ECM in the absence of CD4^+^ T cells (9). The role of CD4 T^+^ cells in activating DC for priming CTL responses during blood stage PbA infection therefore remained to be conclusively defined.

DC are effective antigen presenting cells (APC) that prime T cells for the generation of pathogen-specific adaptive immune responses. The DC compartment is comprised of a heterogeneous array of cell subsets with marked functional specialization (32). Determining the relative contribution of DC subsets to T cell priming is of great importance, especially for the development of new generations of vaccines in which antigen is delivered to a specific DC subset seeking induction of optimal T cell responses. CD8^+^ DC possess specialised machinery to perform antigen cross-presentation to CD8^+^ T cells, which other conventional DC lack (33, 34). However, both CD8^+^ and CD8^−^ DC are capable of antigen presentation to CD4^+^ T cells via MHC class II (35, 36). Consistent with these observations, CD8^+^ DC from mice infected with PbTG were significantly more effective than CD8^−^ DC at presenting OVA antigen to CD8^+^ T cells *in vitro,* whereas both DC subsets were capable of priming CD4 T cells (9). Also, depletion of CD8^+^ DC in PbA infected mice resulted in a sharp decrease in the numbers of activated endogenous CD8^+^ T cells in the spleen, and CD4+ T cell activation was also impaired (37). A similar result was obtained when CD8^+^ DC were depleted in *P. chabaudi* infected mice (38). Another study analysing CD4^+^ T cell responses to *P. chabaudi* antigen showed CD8^+^ DC as superior to CD8^−^ DC at MHC II presentation in the steady state, but the latter were more efficient during infection (39). In the *P. chabaudi* model, mice devoid of CD8^+^ DC developed higher peaks of parasitaemia and more pronounced relapses than their wild-type (WT) counterparts (40). These results suggested an important role for CD8^+^ DC in the development of T cell responses against blood stage malaria parasites.

Here we introduce a CD4^+^ T cell transgenic mouse line specific for a malaria parasite antigen, termed PbT-II, that is cross-reactive with a broad range of malaria parasites including PbA, *P. chabaudi,* and *P. yoelii,* and even the human parasite *P. falciparum.* These cells are also cross-reactive, albeit weakly, to PbA sporozoites. Therefore, PbT-II cells are broadly applicable for the study of CD4^+^ T cell function in multiple malaria models.

## Materials and methods

### Ethics statement

All procedures were performed in strict accordance with the recommendations of the Australian code of practice for the care and use of animals for scientific purposes. The protocols were approved by the Biochemistry & Molecular Biology, Dental Science, Medicine (RMH), Microbiology & Immunology, and Surgery (RMH) Animal Ethics Committee, University of Melbourne (ethic project IDs: 0810527, 0811055, 0911527, 1112347, 1513505).

Mice, mosquitos and parasites

C57BL/6 (B6) mice, MHC I^-/-^ mice (41), IAE^-/-^ mice (MHC II-deficient) (42), Batf3^-/-^ (43), IRF8^-/-^ (44), XCR1 -DTRvenus (45), CD11cDTR (46), CD40^-/-^ (47) and CD40L^-/-^ (48) mice and the transgenic strains OT-II (49), gDT-II (50), PbT-I (16) and PbT-II were used between 6-12 weeks and were bred and maintained at the Department of Microbiology and Immunology. Batf3^-/-^ mice, kindly provided by Kenneth M. Murphy (Washington University), were backcrossed 10 generations to B6 for use in this study. Transgenic T cell lines were crossed to mice expressing GFP ubiquitously (uGFP) or with CD45.1 (Ly5.1) mice to allow for the differentiation of these cells from endogenous T cells after adoptive transfer into recipient Ly5.2 B6 mice. XCR1-DTRvenus and CD11cDTR mice were treated with diphtheria toxin (Calbiochem) as indicated in figure legend. The University of Melbourne. Animals used for the generation of the sporozoites were 4-5 week old male Swiss Webster mice purchased from Monash Animal Services (Melbourne, Victoria, Australia) and maintained at the School of Botany, The University of Melbourne, Australia.

*Anopheles stephensi* mosquitoes (strain STE2/MRA-128 from The Malaria Research and Reference Reagent Resource Center) were reared and infected with PbA as described (51). Sporozoites were dissected from mosquito salivary glands (52), resuspended in cold PBS and irradiated with 2x10^4^ rads using a gamma ^60^Co source. 5 x 10^4^ radiation-attenuated PbA sporozoites (RAS) in 0.2 ml of PBS were intravenously (i.v.) administered to recipient mice.

The rodent malaria lines *Plasmodium berghei* ANKA clone 15cy1 (PbA), *P. berghei* NK65, *P. chabaudi* AS and *P. yoelii* 17XNL were used in this study. Unless otherwise stated, mice were infected i.v. with 10^4^ PbA, *P. berghei* NK65, *P. chabaudi* or *P. yoelii* 17XNL iRBC in 0.2 ml of PBS. The human malaria parasite *P. falciparum* 3D7 was used in *in vitro* analysis.

### Generation of transgenic PbT-II

Transgenic PbT-II mice were generated using the V(D)J segments of the TCRα- and β-genes of a CD4^+^ T cell hybridoma (termed D78) specific for an unidentified blood-stage PbA antigen. This hybridoma was derived from T cells extracted from the spleen of a B6 mouse at day 7 after infection with PbA. 3 x 10^6^ splenocytes from a mouse previously infected with PbA were co-cultured with 5 x 10^5^ conventional DC (extracted from the spleen of FMS-like tyrosine kinase 3 receptor ligand (Flt3-L) treated B6 mice) that were pre-loaded for 2 hours with 2 x 10^6^ PbA schizont lysate as previously described (53) in complete RPMI at 6.5% CO_2_, 37°C. One week later, cultured cells were re-stimulated for a week with fresh DC and PbA schizont lysate. To generate PbA-specific hybridomas, *in vitro* cultured cells were then fused with the BWZ36.GFP fusion partner and exposed to drug selection (54). This led to isolation of the IA^b^-restricted D78 hybridoma from which PbT-II T cell receptor genes were derived. V ✔ and V 🚲 usage by D78 was defined by FACS staining with a panel of V ✔ TCR antibodies and the mouse V 🚲 TCR screening panel (BD Biosciences).

The TCR a region was amplified by PCR from the cDNA of the D78 hybridoma using the forward primer <u>GGATCC</u>AGGAATGGACAAGATTCTG containing a *BamHI* recognition sequence at the 5’ end, designed to bind the 5’ UTR region of V ✔ 2, and the reverse primer C<u>AGATCT</u>CAACTGGACCACAG containing a *BglII* recognition sequence at the 5’ end, specific for the Cα region. Sequencing analysis revealed that the TCR α-chain consisted of V ✔ 2.7, Jα12 and Cα gene segments. The V ✔ 2.7-J ✔ 12-C ✔ segment was cloned into the *BamH*I site of the pES4 cDNA expression vector (55).

TCR Vβ usage was confirmed by PCR on cDNA converted from the RNA of the D78 hybridoma using the primer GAAGATGGTGGGGCTTTCAAGGATC, specific for the Vβ12 gene. Sequencing analysis revealed that the TCR β-chain consisted of Vβ12, Dβ2 and Jβ2.4. The TCR β-chain VDJ segment was amplified by PCR from the genomic DNA of the D78 hybridoma using the forward primer G<u>GATCGAT</u>CACACTTGTTTTCCGTG specific for the Vβ12, incorporating a *Cla*I restriction enzyme site at the 5’ end, and the reverse primer GATCGATCAGCTCACCTAACACGAGGA specific for Jβ2.4 and incorporating a ClaI site at the 5’end and sequenced. The Vβ12-Dβ2-Jβ2.4 segment was found to contain a *Cla*I within its sequence, so a new segment was synthetised in which the *Cla*I site (ATCGAT, coding for the aminoacids Asp and Arg) was changed for ATCGAC (no change in aminoacid sequence). The new segment was cloned into the unique *Cla*I restriction site of the p3A9CβTCR genomic expression vector (49).

Alpha- and beta-chain vector sequences were removed using combined *Cla-*I*/Not-* I and *Apa-*I*/Not-*I restriction enzyme digest, respectively, and coinjected into blastocyts of B6 mice to generate transgenic founder mice.

### Dendritic cell isolation

Dendritic cells were purified from the spleens of mice as previously described (9). Briefly, spleens were finely minced and digested in 1 mg/ml collagenase 3 (Worthington) and 20 μg/ml DNAse I (Roche) under intermittent agitation for 20 min at room temperature. DC-T cell complexes were then disrupted by adding EDTA (pH 7.2) to the digest to a final concentration of 7.9 mM and continuing the incubation for 5 more min. After removing undigested fragments by filtering through a 70 μm mesh, cells were resuspended in 5 ml of 1.077 g/cm^3^ isosmotic nycodenz medium (Nycomed Pharma AS, Oslo, Norway), layered over 5 ml nycodenz medium and centrifuged at 1700 x g at 4^o^C for 12 min. In the experiments to determine MHC restriction by the hybridomas, the light density fraction was collected and DC were negatively enriched by incubation with a cocktail of rat monoclonal anti-CD3 (clone KT3-1.1), anti-Thy-1 (clone T24/31.7), anti-Gr1 (clone RB68C5), anti-CD45R (clone RA36B2) and anti-erythrocyte (clone TER119) antibodies followed by immunomagnetic bead depletion using BioMag goat anti-rat IgG beads (Qiagen).

In the experiments involving analyses of DC subsets, cells obtained after centrifugation in nycodenz medium were stained for CD11c, MHC II, CD8 and CD4 and sorted into (MHC II^hi^ CD11c^hi^) CD8+CD4^−^ (CD8 DC), CD8’CD4+ (CD4 DC) or CD8^−^ CD4^−^ (double-negative, DN DC) using a FACSAria III sorter (BD Biosciences).

DC were resuspended in complete DMEM medium supplemented with 10% foetal calf serum (FCS) before use in functional assays.

Functional assay with hybridomas and IL-2 ELISA

For *in vitro* stimulation assays, DC (5 x 10^4^ per well) were cultured for 1h with titrated amounts of lysed whole blood containing mixed stages of PbA parasites before adding 5 x 10^4^ D78 or PbA-specific MHC I-restricted B4 hybridoma cells [16]. For *ex vivo* experiments, DC were extracted from the spleens of PbA infected mice on day 3 after infection with 10^6^ iRBC.

After culture for 40h at 37^o^C in 6.5% CO_2_, supernatants were collected and concentrations of IL-2 were assessed using the Mouse IL-2 ELISA Ready-Set-Go kit (eBiosciences) following manufacturer’s instructions. Data were represented using logarithmic scales. Because it would not be possible to represent concentration values of “0” in logarithmic scales, all data values were increased by 1. The detection limit of the Mouse IL-2 ELISA Ready-Set-Go kit is 2 pg/ml.

The calculation of the net contribution of DC subsets to MHC II presentation was done by multiplying the mean IL-2 values induced by CD4^+^, CD8^+^ or DN DC in Fig 5B by the relative abundance of the corresponding DC subset in the spleen (in our experiments, MHC II^high^ CD11c^high^ cells in the spleen contained 55.1% CD4+ DC, 13.4% DN DC and 17.7% CD8^+^ DC on average). The numbers obtained from all three DC subsets, representing the IL-2 stimulation corrected to relative DC abundance, were added and the percentage of each individual DC subset in the total number obtain was calculated. This was done for all five DC numbers tested in Fig 5B (3.125x10^3^ to 50x10^3^ DC per well) and the average of the five values obtained for each DC subset was calculated.

### Red Blood Cell coating

Blood was collected from a naive B6 donor. After estimating the concentration of RBC, blood was washed by adding 10 ml DMEM and centrifuging at 3750 rpm for 10 min at 4^o^C (with mild braking). The pellet was then diluted in a solution of DMEM containing 100 mg/ml OVA (Sigma) at a concentration of 10^5^ RBC/μl and incubated at 37^o^C for 30 min. Cells were washed 3 times in PBS before being added to the DC cultures.

To coat RBC with PbA antigen, the concentration of parasites in a blood sample of an infected B6 donor mouse was estimated before lysing by 3 consecutive cycles of freezing/thawing in liquid nitrogen followed by passage through a 30G needle 6 times. The equivalent of 1 parasite/RBC was added to the RBC/OVA solution.

### Generation of bone marrow chimeras

B6 mice were irradiated with two doses of 550 rads 3 hours apart and reconstituted 4h later with 5 x 10^6^ bone marrow cells. These were collected from B6 and CD11c-DTR donor mice and depleted of T cells by coating with antibodies against CD4 (RL172), CD8 (3.168) and Thy1 (J1j) followed by incubation with rabbit complement for 20 min at 37^o^C. One day later, mice were injected intraperitoneally (i.p.) with anti-Thy1 antibody (T24) to deplete residual T cells. Chimeric mice were rested for 8-10 weeks before use, receiving water containing 2.5 g/L neomycin sulphate and 0.94 g/L polymyxin B sulphate for the first 6 weeks after irradiation.

### T cell isolation and *in vivo* proliferation assay

CD4^+^ or CD8^+^ T cells were negatively enriched from the spleens and lymph nodes of PbT-I/uGFP, PbT-II/uGFP, gDT-II/Ly5.1 or OT-II/Ly5.1 transgenic mice as previously described (56) and labelled with CellTrace™ Violet (CTV) or carboxyfluorescein succinimidyl ester (CFSE) following manufacturers instructions (ThermoFisher). 1x10^6^ Purified cells were injected i.v. in 0.2 ml PBS a day before mice were infected with malaria parasites (10^4^, 10^5^ or 10^6^ iRBC or 5x10^4^ RAS, as stated in the figure legend) or HSV (10^5^ pfu HSV). Spleens were harvested at various time points after infection for the analysis of transgenic TCR cell proliferation by flow cytometry.

To deplete endogenous CD4^+^ T cells before adoptive transfer, mice were injected i.v. with 100 μg of anti-CD4 antibody (clone GK1.5) 7 and 4 days prior to the transfer of PbT-II cells.

### *In vitro* proliferation assay

Spleens were isolated from B6 mice that had been infected with 10^6^ PbA iRBC 3 days earlier and DC were enriched by a 1.077 g/cm^3^ nycodenz density centrifugation followed by negative selection. 2x10^5^ DC and 10^5^ CTV-labelled PbT-II cells were incubated with titrated amounts of *P. falciparum-* or PbA-iRBC for 3 days. PbT-II proliferation was assessed by flow cytometry.

### Flow cytometry

Cells were labelled with monoclonal antibodies specific for CD8 (clone 53-6.7), CD4 (RM 4-5), Thy1.2 (30-H12), MHC II (M5/114.15.2), CD11c (N418), CD45.1 (A20), Vα2 (B20.1), Vβ12 (MR11-1), Vα8.3 (B21.14), νβ10 (B21.5) or CD69 (H1.2F3), Sirpα (P84), Tbet (4B10), Bcl6 (K112-91), Foxp3 (FJK-165), GATA3 (L50-823), RORγt (Q31- 378). Dead cells were excluded by propidium iodide staining. Cells were analyzed by flow cytometry on a FACS Canto or Fortessa (BD Biosciences), using the Flowjo software (Tree Star Inc.).

For intracellular staining, cells were permeabilised using the Transcription Factor Buffer Set (BD Biosciences) following manufacturer’s instructions. Dead cells were excluded using LIVE/DEAD fixable dead cell stain (Thermofisher).

For the determination of parasitaemia, 1-2 μl blood were collected from the tail vein, diluted in FACS buffer (containing 1% w/v BSA and 5mM EDTA) and incubated at 37^o^C, 6.5% CO_2_ with 5μg/ml Hoechst 33258 (Thermofischer) for 1h before running on a Fortessa. Blood from naïve mice was used as negative control (background stain usually ranged between 0.10-0.14%)

### Generation and monitoring of ECM

Mice infected with blood-stage PbA were monitored daily for the development of ECM. Mice were considered to have ECM when showing signs of neurological symptoms such as ataxia and paralysis, evaluated as the inability of mice to self-right.

### Detection of *P. chabaudi*-specific antibody

Nunc-Immuno™ MicroWell™ 96-well ELISA Plates (300μL round-bottom) were coated overnight at 4^o^C with 50μL PBS containing *P. chabaudi* blood-stage lysate (equivalent to 6.6x10^5^ parasites/well). Unbound antigen was washed away by soaking the plates in 0.05% tween 20/PBS (i.e. wash) ×4 times. The wells were blocked with 50μL of 5% skim milk/PBS for 15-30min. Plasma samples were serially diluted in 5% skim milk/PBS and incubated overnight at 4^o^C. The wells were washed ×6 with 0.05% tween 20/PBS and incubated overnight at 4^o^C with 50μL of 5% skim milk/PBS containing antimouse IgG-HRP (1:10,000) to assess the total IgG responses. The wells were washed ×6 with 0.05% tween 20/PBS and HRP was detected by adding 50μL ABTS substrate in the wells and incubating for 1.5-2h at RT. Optical density at 405nm (OD_405nm_) and the background at 492nm (OD_492nm_) was determined using an ELISA plate reader. The endpoint titers were calculated by using cut-off values determined as 2×SD above average OD_405nm-492nm_ values of control wells containing no plasma.

*P. chabaudi* antigen used to coat ELISA plates was prepared as follows: Blood from *P. chabaudi*-infected mice was incubated with 0.05% saponin for 3 min at RT to release parasites from RBC. Parasites were then broken up by snap freezing in liquid nitrogen and thawing 3 times, followed by 6 passages through a 30G needle.

### Statistics

Data were log transformed for conversion into a normal distribution and then analysed using parametric statistical tests such as t-test for the comparison of 2 groups or ANOVA followed by Tukey’s Multiple Comparison Test for simultaneous comparison of multiple groups. *P*-values <0.05 (*), <0.01 (**) or 0.001 (***) were considered statistically significant.

## Results

### Generation of an MHC Il-restricted T cell hybridoma specific for PbA

To study the CD4^+^ T cell response to blood stage malaria, we developed an MHC II- restricted T cell hybridoma specific for PbA. This hybridoma, termed D78, was derived by fusing the immortalized cell line BWZ36.GFP (57) to CD4^+^ T cells isolated from the spleen of a B6 mouse infected with blood stage PbA. To assess MHC restriction of D78, DC deficient in MHC I or MHC II molecules were pre-incubated with lysate of PbA-infected red blood cells (iRBC) and then tested for stimulatory capacity (Fig 1). Whereas both WT and MHC I knockout (KO) DC were able to efficiently stimulate this hybridoma, IA^b^ deficient (MHC II KO) DC were non-stimulatory. In contrast, MHC II KO DC were able to stimulate a PbA-specific MHC I-restricted hybridoma (16), showing these DC were functional (Fig S1A). Furthermore, D78 responded efficiently to PbA iRBC lysate, but failed to respond to DC activated by non-specific stimuli such as LPS, polyI:C or CpG, thus excluding self-reactivity or reactivity to foetal calf serum proteins (Fig S1B). Together, these findings indicated that the D78 hybridoma was MHC II-restricted and specific for malaria antigen.

**Fig 1.**
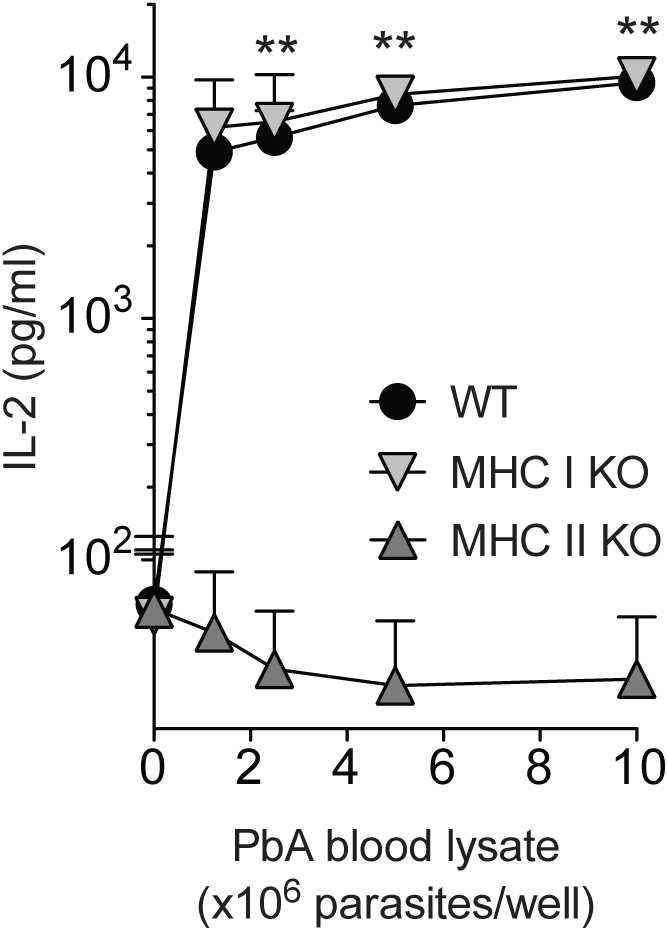
MHC II restriction of the D78 hybridoma. Dendritic cells were enriched from the spleens of WT B6 (filled circle), MHC I deficient (inverted triangle) or MHC II deficient (upright triangle) mice and cultured for 1h with titrated amounts of lysed blood-stage PbA iRBC before adding 5x10^4^ D78 hybridoma cells. 40h later, IL-2 concentrations in the supernatants were measured by ELISA. Data points denote mean of IL-2 concentration and error bars represent SEM. Data were pooled from 2 independent experiments. Statistical analysis was performed on log-transformed data using unpaired t tests. Asterisks denote significant differences between WT and MHC II deficient DC (^**^, P<0.01). No statistical differences were found between WT and MHC I deficient DC.

### Generation of a PbA-specific TCR transgenic line

To enable monitoring of CD4^+^ T cell immunity to PbA *in vivo,* an MHC II-restricted T cell receptor (TCR) transgenic mouse line was developed using the TCR alpha and beta chains expressed by D78. This line has been termed PbT-II, consistent with our nomenclature for MHC I and II restricted OVA-specific lines OT-I and OT-II and our recently reported MHC I-restricted PbA-specific line specific, termed PbT-I (16). Analysis of the spleen and inguinal lymph node (iLN) of PbT-II mice revealed skewing towards CD4+ T cells as well as efficient expression of the Vα2 and Vβ12 transgenes derived from D78 (Fig 2 and S1D). Total cellularity in the spleen of PbT-II mice was similar to WT mice, with CD4^+^ T cells largely compensating for reduced numbers of CD8^+^ T cells in the former (Fig S1E). Some reduction in the total cellularity of the iLN was evident, but the reason for this is unclear. Of note, increased numbers of CD4^−^ CD8^−^ (double-negative (DN)) T cells were also found in the spleen and iLN of PbT-II mice. In the thymus, PbT-II mice showed an increased number and proportion of single positive CD4^+^ T cells and double negative T cells (Fig S1C and S1E Fig). Similar to previously described TCR transgenic mice, total cellularity was reduced in the thymus, indicative of efficient positive selection (16, 58).

**Fig 2.**
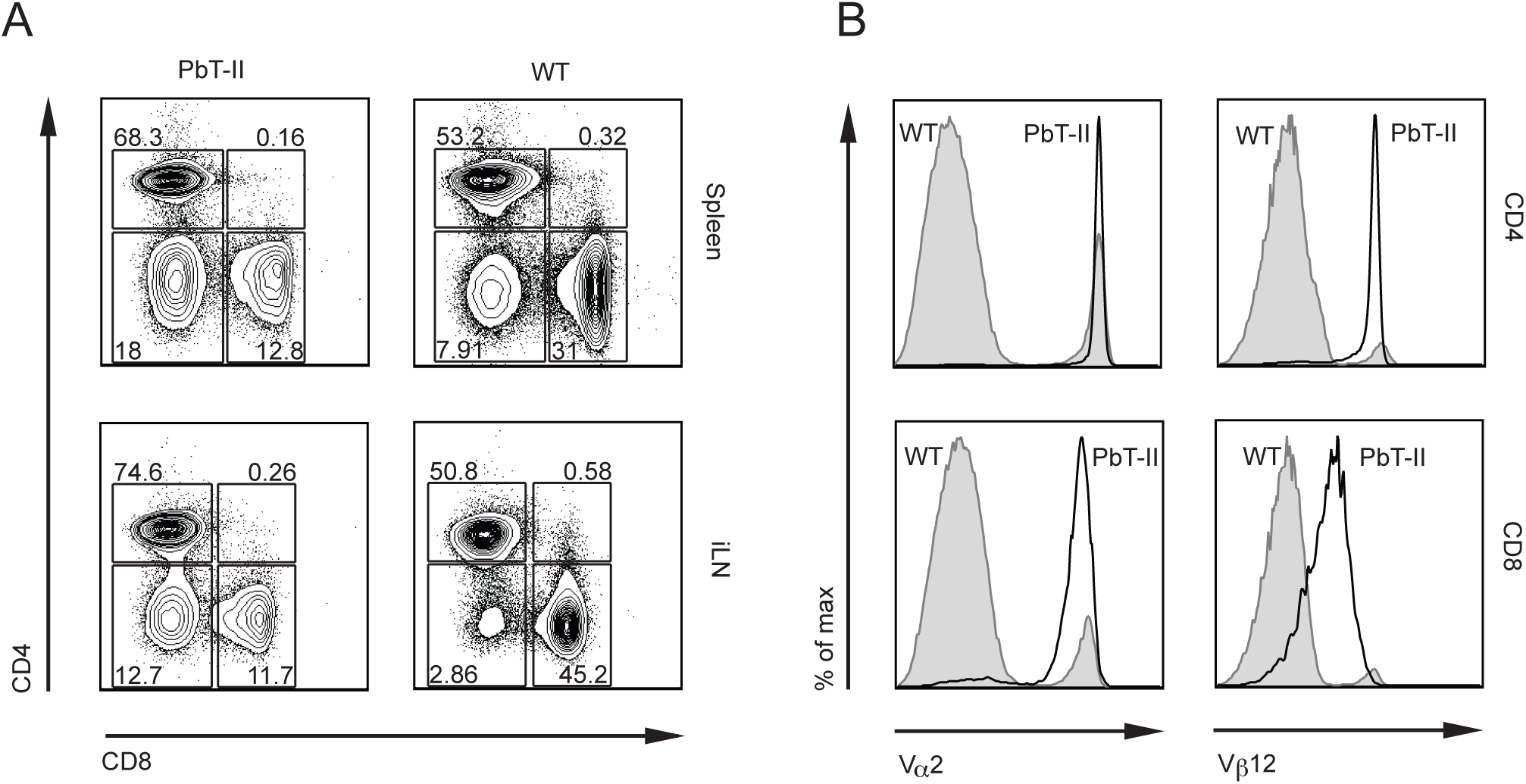
Characterization of T cells from the spleen and lymph nodes of PbT-II mice. Cells were harvested from the spleen and the lymph nodes of PbT-II transgenic or control B6 (WT) mice. FACS analysis was performed to characterize the expression of CD4, CD8 and the transgenic TCR alpha (V ✔ 2) and beta (V 🚲 12) chains. (A) Representative plots showing the proportions of CD4^+^ versus CD8^+^ Thy1.2^+^ cells in the spleen and inguinal lymph node of PbT-II and WT mice. (B) Representative histograms showing the expression of the transgenic TCR V ✔ 2 and V 🚲 12 chains on the CD4 or CD8 single positive cells from the spleen. Data are representative for 2 independent experiments.

### PbT-II cells respond to multiple *Plasmodium* species and life-cycle stages

PbT-II mice were generated without knowledge of their specific peptide antigens. To further characterise the specificity of this line, we tested responsiveness to different species of malaria parasites. CellTrace™ Violet (CTV)-labelled PbT-II cells were adoptively transferred into B6 mice one day before infection with iRBC from either PbA, *P. berghei* NK65, *P. chabaudi* or *P. yoelii* 17XNL. Several days later, PbT-II proliferation was assessed in the spleen, revealing reactivity to all *Plasmodium* species (Fig 3A). This result prompted us to determine whether PbT-II cells also responded to the human malaria parasite *P. falciparum. In vitro* culture of PbT-II cells with DC and blood-stage *P. falciparum* lysate resulted in PbT-II proliferation comparable to that seen in response to PbA (Fig 3B), indicating that the antigen recognised by PbT-II cells is conserved in multiple *Plasmodium* species.

**Fig 3.**
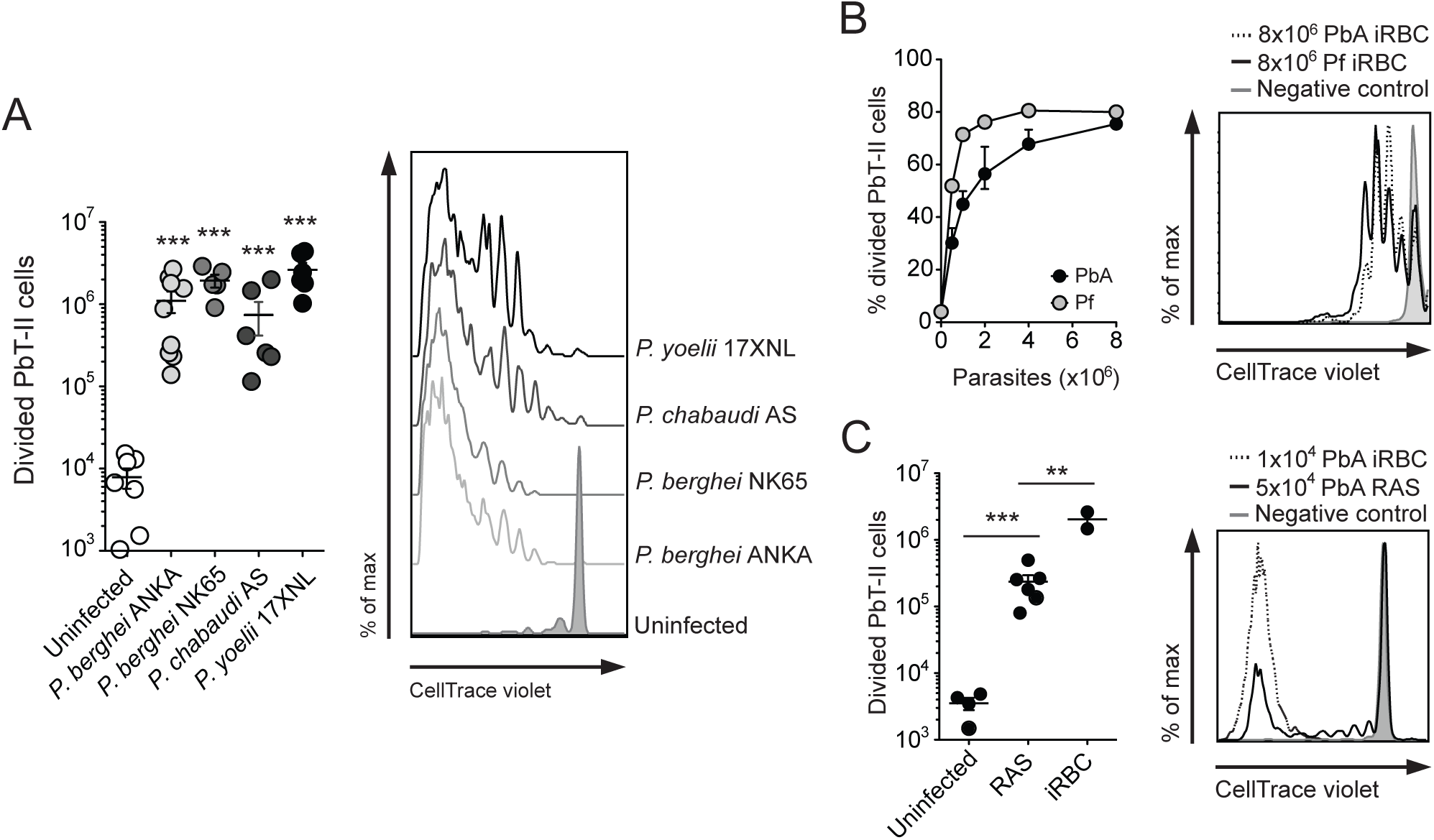
PbT-II cells are cross-reactive with multiple *Plasmodium* species and with liver stage PbA. (A) Cross-reactivity with murine malaria parasites. 10^6^ CTV-labelled PbT-II cells were transferred into B6 recipients one day before infection with 10^4^ iRBC from *Plasmodium berghei* ANKA (PbA), *P. berghei* NK65, *P. chabaudi* AS, *P. yoelii* 17XNL or nothing. 5 days after *P. berghei* ANKA and *P. berghei* NK65 infection, or 7 days after *P. chabaudi* AS and *P. yoelii* 17XNL infection, mice were killed and PbT-II proliferation was assessed in the spleen. *Left.* Number of proliferating cells. *Right.* Representative histograms showing PbT-II proliferation. Data are representative for 2 independent experiments. Data were log transformed and analysed using One Way ANOVA and Dunnet’s Multiple comparisons test. Asterisks denote statistical differences with the uninfected group (^***^, P<0.001) (B) Cross-reactivity with *P. falciparum.* 2x10^5^ Splenic DC were cultured with 10^5^ CFSE-coated PbT-II cells and titrated amounts of *P. falciparum* (Pf)- or PbA-iRBC for 3 days. PbT-II proliferation was assessed by flow cytometry. *Left.* Percentages of proliferating PbT-II cells to Pf or PbA antigen. *Right.* Representative histogram showing PbT-II proliferation to Pf-iRBC (solid black line), PbA iRBC (dotted line) and a negative control containing no antigen (tinted grey line). (C) Cross-reactivity with liver stage PbA parasites. 10^6^ CTV-labelled PbT-II cells were transferred into recipient B6 mice one day before infection with 5x10^4^ *P. berghei* ANKA radiation attenuated sporozoites (RAS) or 10^4^ PbA iRBC. 6 days later mice were killed and PbT-II proliferation was assessed in the spleen. *Left.* Number of proliferating cells. *Right.* Representative histograms showing PbT-II proliferation. The tinted line represents uninfected control. The solid and dotted black lines show PbT-II cell proliferation in mice infected with PbA RAS or iRBC respectively. Data were pooled from 2 independent experiments. Data were log transformed and analysed using One Way ANOVA and Tukey’s Multiple comparisons test (^**^, P<0.01; ^***^, P<0.001)

We then asked whether the antigen recognized by PbT-II cells was also expressed by preerythrocytic stage PbA parasites. To examine this, CTV-labelled PbT-II cells were adoptively transferred into B6 mice that were then infected with radiation-attenuated sporozoites (RAS) that do not develop into blood stage infection. This revealed significant, though weak proliferation of PbT-II cells 6 days later, indicating some reactivity to the preerythrocytic stage of PbA (Fig 3C).

Because the PbT-II cells proliferated in every assay we performed, we wanted to confirm *in vivo* that such responsiveness was due to the recognition of a malaria parasite-derived epitope, and not to non-specific inflammatory signals. We transferred gDT-II cells, which are CD4^+^ T cells specific for HSV, and PbT-II cells into B6 mice that were intravenously infected with 10^5^ HSV plaque-forming units (pfu) one day later. Whereas gDT-II cells proliferated extensively by day 10 after infection, PbT-II cells remained undivided (Fig S1F).

Taken together, these results revealed a broad responsiveness of PbT-II cells to malaria parasites, including different life-cycle stages and species, showcasing the potential for these T cells as tools to broadly study CD4^+^ T cell responses in malaria.

### PbT-II cells can provide immunity to_*P*. *chabaudi* infection

Both B cells and helper T cells are essential for the control of *P. chabaudi* parasitaemia (59, 60) and protection is mainly achieved by parasite-specific T cell-dependent antibodies (17). This model was therefore ideally suited to test whether PbT-II cells have the capacity to provide protective immunity to malaria. We transferred PbT-II cells into CD40L- deficient mice, in which endogenous CD4^+^ T cells are unable to provide help to B cells (47), and infected these mice with 10^4^ *P. chabaudi* iRBC. CD40L-deficient mice lacking PbT-II cells succumbed to infection during the peak of parasitaemia on day 9 p.i., or shortly thereafter, whereas those adoptively transferred with PbT-II cells survived >30 days (Figure 4A, S2). These latter mice developed a second peak of parasitaemia on day 19 p.i. that occurred earlier and was higher than that of wild-type (WT) mice (Fig 4B and S2). Although parasitaemia was still detectable on day 26 after infection, CD40L-deficient mice that received PbT-II cells had recovered from disease symptoms, as revealed by increased body weight to pre-infection levels (Fig S2), and were sacrificed >50 days after infection without signs of disease. Analysis of the plasma of these mice on day 9 after infection also revealed increased levels of *P. chabaudi*-specific IgG in CD40L-deficient mice that received PbT-II cells compared to those that did not receive cells (Fig 4C).

**Fig 4.**
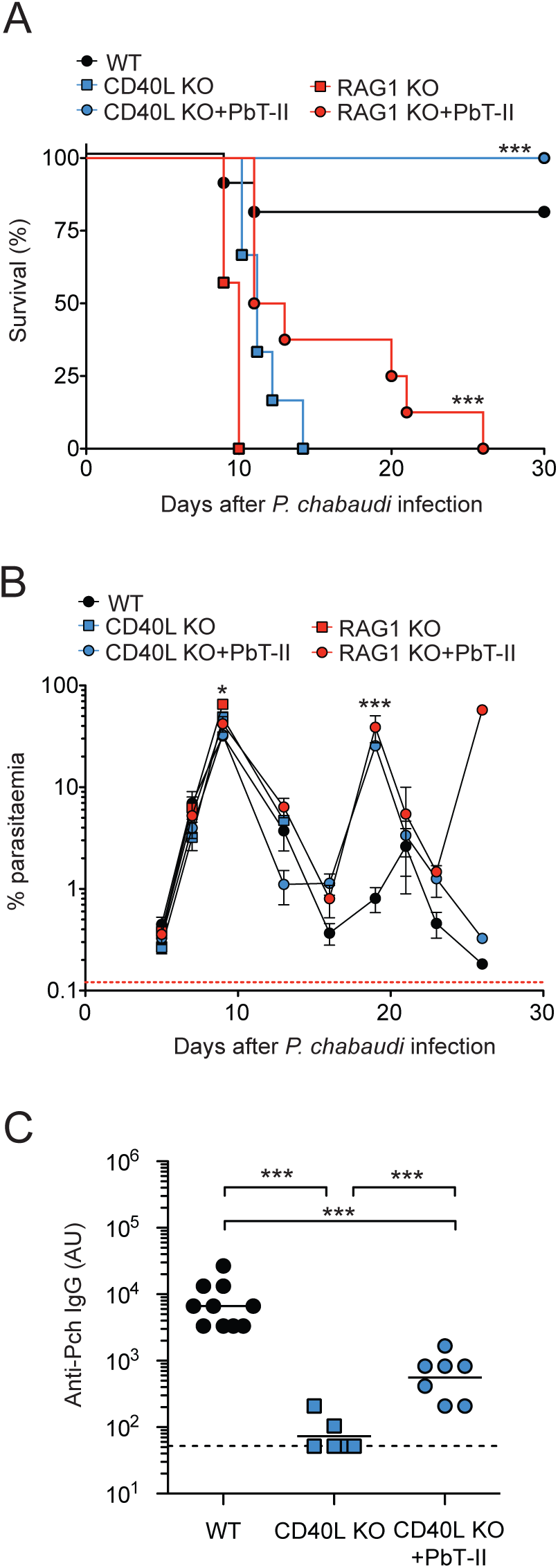
PbT-II cells can elicit immunity to *P. chabaudi.* infection. CD40L-deficient mice (CD40L KO), RAG-deficient mice (RAG KO) and WT control mice received either 5x10^5^ PbT-II cells or no cells one day before infection with 10^4^ *P. chabaudi* iRBC. (A) Survival of WT mice (n=10), CD40L-deficient mice with or without PbT-II cells (n=7 and 6 respectively) and RAG-deficient mice with or without PbT-II cells (n=8 and 7 respectively). Data were analysed using logrank test. Asterisks denote statistical difference between CD40L KO vs CD40L KO+PbT-II cells and RAG KO vs RAG KO+PbT-II cells respectively (^***^, P<0.001). (B) Course of parasitaemia after *P. chabaudi* infection. Blood samples were examined by FACS. The dotted red line represents background staining in uninfected mice. Data were log transformed and analysed using one way ANOVA and Tukey’s Multiple Comparison test. A statistical difference was found on day 9 between RAG KO and RAG KO+PbT-II (^*^, P<0.05). No difference was found between CD40L KO and CD40L KO+PbT-II mice on day 9. Statistically significant differences between both CD40L KO+PbT-II or Rag KO+PbT-II mice and WT mice was found on day 19 (^***^, P<0.001). (C) Parasite-specific IgG antibodies in CD40L KO mice. *P.* chabaudi-specific IgG end-point titers were determined on day 9 after infection using ELISA. Cut-off values were determined as 2×SD above average OD_405nm-492nm_ values of control wells containing no plasma. Data were log transformed and analysed using a One Way ANOVA and Tuckey’s Multiple Comparison Test (^***^, P<0.001). The dotted line represents the limit of detection of the assay.

The protective effect of PbT-II cells for *P. chabaudi* infection was not limited to the production of antibodies, as adoptive transfer of PbT-II cells into RAG1-deficient mice, which are devoid of T cells and B cells, also resulted in a significantly prolonged survival (Fig 4A). The first peak of parasitaemia, on day 9 after infection, was significantly higher in RAG1-deficient mice that did not receive PbT-II cells than in WT mice and RAG1- deficient mice that received PbT-II cells (Fig 4B and S2). Unlike as seen for CD40L-deficient mice, however, PbT-II cells did not enable RAG1-deficient mice to fully control parasitaemia and these mice eventually succumbed to infection (Fig 4A, B and S2).

These results clearly show PbT-II cells promoted immunity against *P. chabaudi* infection via antibody production and antibody-independent mechanisms that contributed to the control of parasitaemia.

### CD8^+^ Dendritic Cells are the main antigen presenting cells for PbT-II cells

Previous reports using DC from infected mice *ex vivo* have implicated both CD8^+^ and CD8^−^ DC in MHC II-restricted antigen presentation during blood-stage malaria infection (9, 39, 61). To examine the capacity of conventional DC (cDC) subsets to present malaria antigens to CD4+ T cells, we enriched cDC (defined as CD11c^high^ MHC class II^high^ cells) from the spleens of naïve mice and subdivided them into three subsets: CD8^+^CD4^−^, CD8^−^CD4^+^ and CD8^−^CD4^−^ DC (CD8^+^, CD4^+^ and DN DC respectively) (62). CD8^+^ DC also express XCR1 (45), Clec9A (63), CD24 and DEC205 (62), and lack expression of Sirpα (Fig S3A). CD4^+^ DC are the most abundant of the three subtypes (making up around 55% of all cDC), and express CD11b (62) and Sirpα (Fig S3A). DN DC are a heterogeneous group of DC in which most cells express Sirpα and CD11b, but a small group lack Sirpα (Fig S3A), likely cDC precursors(64). We incubated purified DC subtypes with PbA iRBC and the D78 hybridoma *in vitro* and assessed the hybridoma responses (Fig 5A). While all three DC subsets were capable of stimulating D78, their efficacy varied considerably: CD8^+^ DC induced the strongest responses; DN DC had an intermediate stimulatory capacity and CD4^+^ DC were significantly less efficient than any of the other subtypes. A similar result was obtained when the three cDC subsets were purified from the spleens of B6 mice that had been infected with PbA 3 days earlier (Fig 5B). In this case, the stimulation exerted by DN DC and CD4^+^ DC was comparable and significantly lower than that of CD8^+^ DC.

It was possible that the observed superiority of CD8^+^ DC was due to the characteristics of the specific peptide recognised by the D78 hybridoma, and not generalizable to other peptides. To clarify this point, we sought to test the capacity of cDC subsets at presenting OVA to the OT-II hybridoma *in vitro.* Consistent with previous studies (34), incubation of CD4^+^, CD8^+^ and DN DC with soluble OVA demonstrated a comparable capacity of all these DC to stimulate the OT-II hybridoma (Fig 5C). However, because the form of the antigen influences the effectiveness of its capture and processing by DC (34), and the malaria parasite antigen used to assess D78 responses *in vitro* in Fig 5A was cell-associated (i.e. from iRBC), we coated RBC from a naive B6 mouse with OVA in an attempt to provide this antigen to DC in a comparable, cell-associated fashion. Similar to the dominance of CD8^+^ DC in presentation of iRBC malaria antigen to D78, these DC were the most efficient stimulators of the OT-II hybridoma when OVA-coated RBC were used (Fig 5D). CD4+ DC were the least efficient DC subtype and DN DC exhibited an intermediate stimulatory capacity.

**Fig 5.**
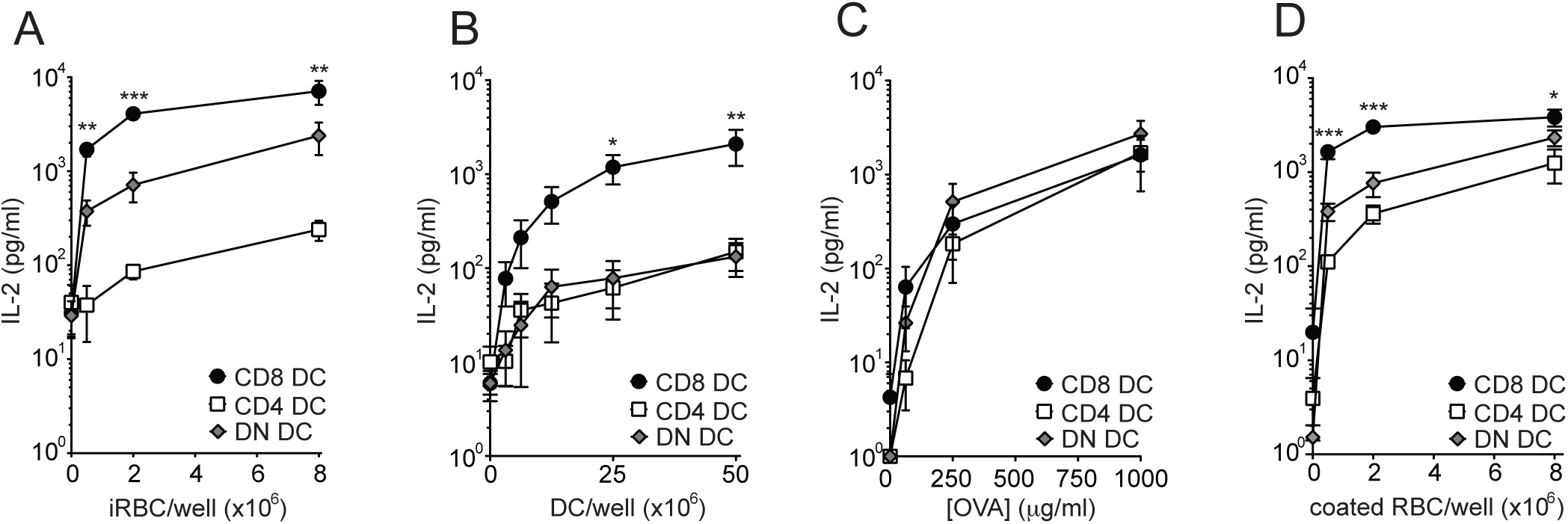
CD8^+^ DC are more efficient than other DC subtypes at stimulating D78 hybridoma cells *in vitro* and *ex vivo.* (A) D78 stimulation by DC subtypes *in vitro.* 5x10^4^ sorted CD4^+^, CD8^+^ or DN DC from naive B6 mice were pre-incubated for 1h with titrated amounts of PbA iRBC. 5x10^4^ D78 hybridoma cells were then added to the culture and incubated for a further 40h before supernatants were collected and IL-2 concentrations measured by ELISA. (B) D78 stimulation by DC subtypes *ex vivo.* 5x10^4^ sorted CD4^+^, CD8^+^ or DN DC from the spleens of B6 mice infected with 10^6^ PbA iRBC 3 days earlier were incubated with 5x10^4^ D78 hybridoma cells for 40h before supernatants were collected and IL-2 concentrations measured by ELISA. (C) Presentation of soluble OVA antigen to the OT-II hybridoma. 5x10^4^ sorted CD4^+^, CD8^+^ or DN DC from naive B6 mice were preincubated for 1h with titrated amounts of soluble OVA. 5x10^4^ OT-II hybridoma cells were then added to the culture and incubated for a further 40h before supernatants were collected and IL-2 concentrations measured by ELISA. (D) CD8^+^ DC are more efficient than other DC subtypes at presenting RBC-associated antigen to the OT-II hybridoma *in vitro.* 5x10^4^ sorted CD4^+^, CD8^+^ or DN DC from naive B6 mice were pre-incubated for 1h with OVA-coated RBC from a naive B6 mouse. 5x10^4^ OT-II hybridoma cells were then added to the culture and incubated for a further 40h before supernatants were collected for assessment of IL-2 concentrations by ELISA. Data were pooled from 3 independent experiments in AC and from 4 independent experiments in D. Data were log-transformed and statistically analysed using One-way ANOVA (^*^, P<0.05; ^**^, P<0.01; ^***^, P<0.001).

Together, these results indicate that CD8^+^ DC have the highest capacity for stimulation of PbT-II T cells and suggest that this DC subset is particularly efficient at presenting MHC II-restricted antigens associated with RBC.

To validate the role of CD8^+^ DC *in vivo,* we then took advantage of the PbT-II TCR transgenic mice, which enabled us to follow the response of adoptively transferred PbT-II T cells in mice lacking DC. Although DC are efficient APC capable of activating CD4^+^ T cells, other cells can present antigen via MHC II and can therefore contribute to CD4^+^ T cell priming (65). To assess the contribution of DC in the priming of PbT-II cells during PbA infection, we first reconstituted lethally-irradiated B6 mice with bone marrow from CD11c-DTR mice. In these chimeras, DC could be depleted by the administration of diphtheria toxin (DT) (46). Consistent with previous reports (9, 66), DT treatment of CD11cDTR->B6 chimeras during PbA infection resulted in the complete abrogation of PbT-II cell proliferation (Figs 6A and E), demonstrating an essential role of DC in CD4+ T cell priming during blood stage infection. A similar result was obtained when DC were removed after *P. chabaudi* infection (Fig S3B).

**Fig 6.**
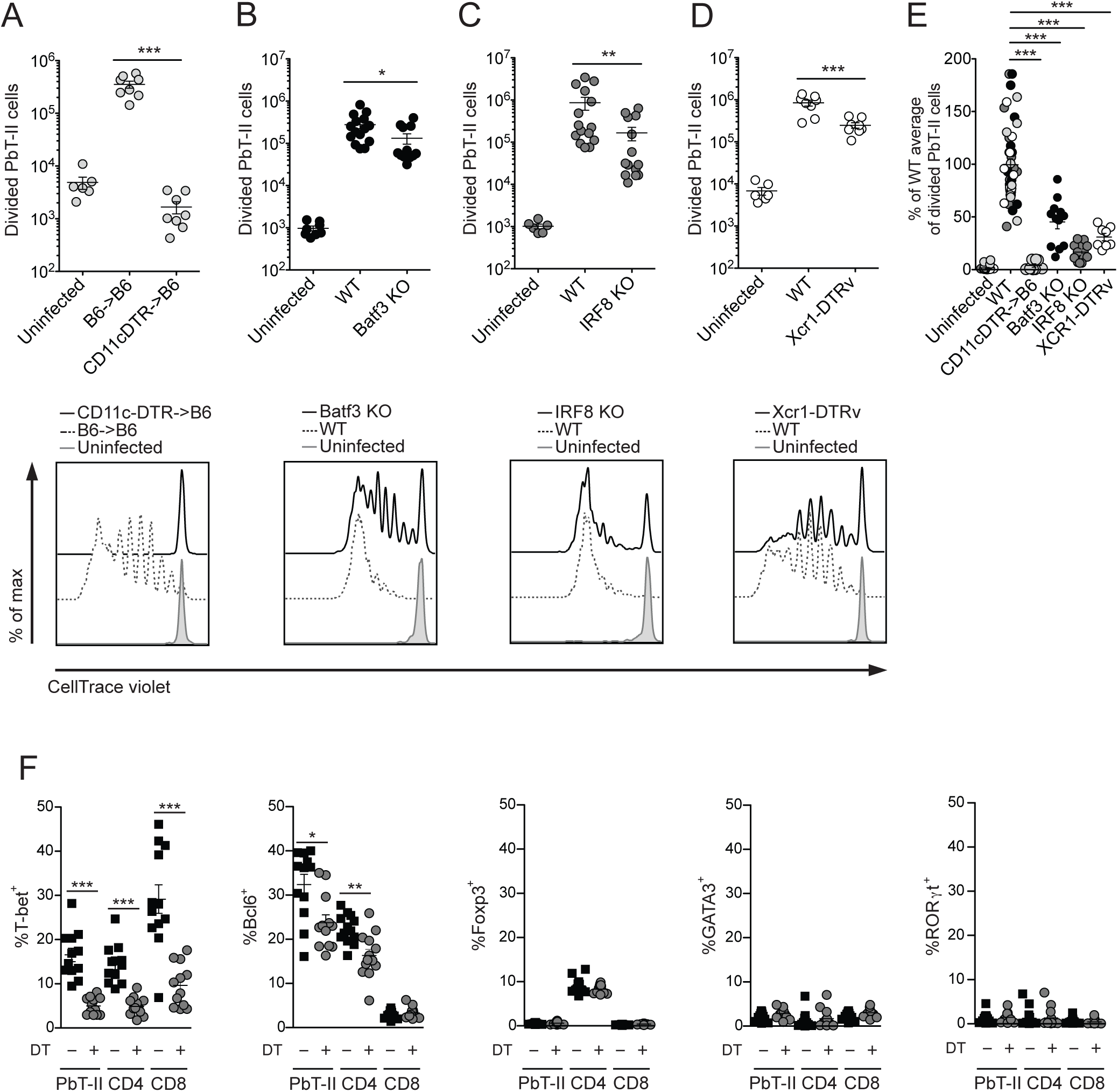
PbT-II priming during blood stage PbA infection is exclusively done by DC, among which the CD8^+^ subset is the major contributor. 10^6^ CTV-labelled PbT-II cells were adoptively transferred into different DC deficient recipient mice or WT controls, which were infected with 10^4^ PbA iRBC 1-2 days later. Proliferation of PbT-II cells was measured in the spleen on day 5 after PbA infection. Total divided cells in the spleen (upper graph) and the corresponding representative histograms for proliferation (below) are shown in each subfigure. The different groups in the upper graph are represented in the histograms below as solid black lines (DC deficient mice), dotted lines (WT control mice) and tinted grey lines (uninfected mice). (A) PbT-II proliferation in DT-treated B6->B6 or CD11c- DTR->B6 chimeras. Mice were treated with 100 μg DT i.p. on days -1, 1 and 3 after infection. (B) PbT-II proliferation in WT or Batf3 KO mice. (C) PbT-II proliferation in WT or IRF8 KO mice. (D) PbT-II proliferation in DT-treated WT or XCRl-DTRvenus mice. Mice were treated with 500 μg DT i.p. on days -1, 0, 1 and 2 after infection. Data were pooled from 2 independent experiments in A and D, and 3 independent experiments in B and C. For statistical analysis, data were log transformed and analysed using one way ANOVA followed by Tukey’s Multiple comparisons test (n.s.=not significant; ^*^, P<0.05, ^**^, P<0.01, ^***^, P<0.001). (E) Percentages of PbT-II proliferation in DC deficient mice compared with their WT counterparts. Data derived from A-D was normalized to facilitate comparisons between individual experiments. First, the average of divided PbT-II cells in WT mice in each experiment was calculated. That number was then used to calculate the percentage of PbT-II divided cells in both KO mice (CD11cDTR->B6, light grey circles; Batf3 KO, black circles; IRF8 KO, dark grey circles; XCR1-DTRvenus mice, white circles) or their respective WT counterparts (whose normalised proliferation would therefore average 100%). Data in E were statistically compared using Kruskal-Wallis tests followed by Dunn’s multiple comparisons test for differences between the individual groups (^***^, P<0.001). (F) Transcription factor expression in splenic PbT-II cells and endogenous CD4^+^ and CD8^+^ T cells primed in the presence or absence of CD8^+^ DC. XCR1-DTRvenus mice received 5x10^5^ PbT-II/uGFP cells and were infected with 10^4^ PbA iRBC one day later. These mice were treated with either DT (500μg/dose) or PBS on days -1, 0, 1, 2, 4 and 6 p.i. and killed on day 7. Data points represent individual mice that were pooled from 2 independent experiments, log transformed and statistically analysed using an unpaired t test (^*^, P<0.05, ^**^, P<0.01, ^***^, P<0.001). Data are represented as mean±SEM.

To assess the role of individual DC subsets in the PbT-II response, we examined reactivity in mice deficient in subsets of DC due to the lack of specific transcription factors, namely Batf3 (43) or IRF8 (44). Batf3 deficient mice contain all DC subsets except CD8+ DC and their migratory equivalents, the CD103^+^ DC, and have been shown to sustain low CD8^+^ T cell proliferation after PbA infection (67),(43). However, on a B6 background, some CD8^+^ DC can be found in the LN of Batf3-deficient mice (68) and precursors capable of MHC II-restricted presentation but not cross-presentation are found in the spleen (69). To examine the PbT-II response in Batf3-deficient mice, CTV-labelled PbT-II cells were adoptively transferred one day before infection with 10^4^ PbA iRBC. PbT-II proliferation was then assessed 5 days later (Figs 6B and E). This showed that the total number of proliferating PbT-II cells was reduced in Batf3 deficient mice (Fig 6B), and pair-wise comparisons showed a 50% reduction relative to WT controls (Fig 6E). These findings indicated that PbT-II cells were heavily dependent upon mature CD8+ DC for antigen presentation, but suggested some participation by other DC, possibly the less mature form of CD8^+^ DC present in Batf3-deficient spleens (incapable of cross-presentation) or an alternative CD8^-^ DC subset.

IRF8-deficient mice are profoundly deficient in CD8^+^ DC, CD103^+^ DC and pDC, but contain other conventional DC subsets, so these mice represent an ideal host for assessing the role of CD8^−^ conventional DC in antigen presentation. CTV-labelled PbT-II cells were therefore adoptively transferred into IRF8-deficient mice 1 day before infection with PbA iRBC and T cell proliferation assessed 5 days later (Figs 6C and E). PbT-II proliferation was severely reduced in these mice (Fig 6C), amounting to about 20% of WT responses (Fig 6E), further supporting the major role for CD8^+^ DC in PbT-II responses.

Lack of IRF8 is known to result in immune defects in addition to the absence of CD8^+^ DC, e.g. CD8^−^ DC, although in normal numbers, are hypo-responsive to microbial stimulation (70). Also, IRF8 deficient mice show an excessive production of granulocytes resulting in splenomegaly (44), which could have hindered proper priming and proliferation of PbT-II cells. Thus, to further explore the importance of CD8+ DC in PbT-II priming to PbA, we assessed PbT-II proliferation in DT-treated XCR1-DTRvenus mice. XCR1 is a chemokine receptor expressed by mature CD8^+^ DC and by their migratory CD103^+^ counterparts, found in lymph nodes. DT treatment of XCR1-DTRvenus mice, which express the DT receptor under the XCR1 promoter, results in the elimination of CD8^+^ DC from the spleen (45). Consistent with the results obtained using IRF8-deficient mice infected with blood stage PbA, PbT-II proliferation was significantly reduced in the spleen of DT-treated XCR1- DTRvenus mice, amounting to about 30% of WT controls (Figs 6D and E).

Together, these data demonstrated an essential role for DC in the generation of PbT-II cell responses to blood stage malaria parasites and argued that the CD8^+^ DC subtype played a dominant role.

### CD8^+^ DC influence Th1 and Tfh induction

DC subsets can influence the expression of transcription factors by T cells and drive their differentiation into particular T helper phenotypes (35, 36, 71-73). To address the role of CD8^+^ DC on the differentiation of PbT-II cells after PbA infection, XCR1-DTRvenus mice were adoptively transferred with PbT-II cells and then infected with 10^4^ PbA iRBC. These mice were either treated with DT throughout the infection to remove CD8^+^ DC or given PBS, maintaining this DC subset. On day 7, the expression of transcription factors that define Th1, Th2, Th17, Tfh and Treg subsets (i.e. Tbet, GATA3, RORγt, Bcl6 and Foxp3 respectively) was assessed in PbT-II cells revealing induction of Tbet^+^ and Bcl6^+^ PbT-II cells, which was impaired upon depletion of CD8^+^ DC (Fig 6F and S3C). No expression of GATA3, RORγt or Foxp3 was detected in PbT-II cells, even in the presence of CD8^+^ DC. A similar dependence on CD8^+^ DC for Th1 and Tfh cell development was observed for endogenous CD4^+^ T cells (Fig 6F). Of note, and agreeing with a previous study (73), the proportion of Tbet^+^ CD8^+^ T cells was also markedly reduced in mice depleted of CD8^+^ DC. These results suggested a major role for CD8^+^ DC in promoting Tbet expression in CD4 and CD8 T cells, and a lower, but significant contribution to the generation of Tfh cells.

### CD8^+^ DC function early after infection is essential for ECM development

*In vitro* experiments involving the use of transgenic parasites expressing model antigens previously showed that CD8^+^ T cell priming during blood stage malaria parasite infection is efficiently performed by CD8^+^ DC, but only poorly by other DC subsets (9). To confirm this point *in vivo,* we transferred CD8^+^ T cells from the PbT-I transgenic line into Batf3- deficient mice or XCR1-DTRvenus mice and then infected these mice with PbA. XCR1- DTRvenus mice were treated with DT from days -1 to 2 after infection to specifically remove CD8^+^ DC. PbT-I proliferation was severely decreased in the absence of CD8^+^ DC in both cases (Fig 7A and B), confirming the important role of this DC subtype in CD8^+^ T cell activation during blood stage PbA infection.

**Fig 7.**
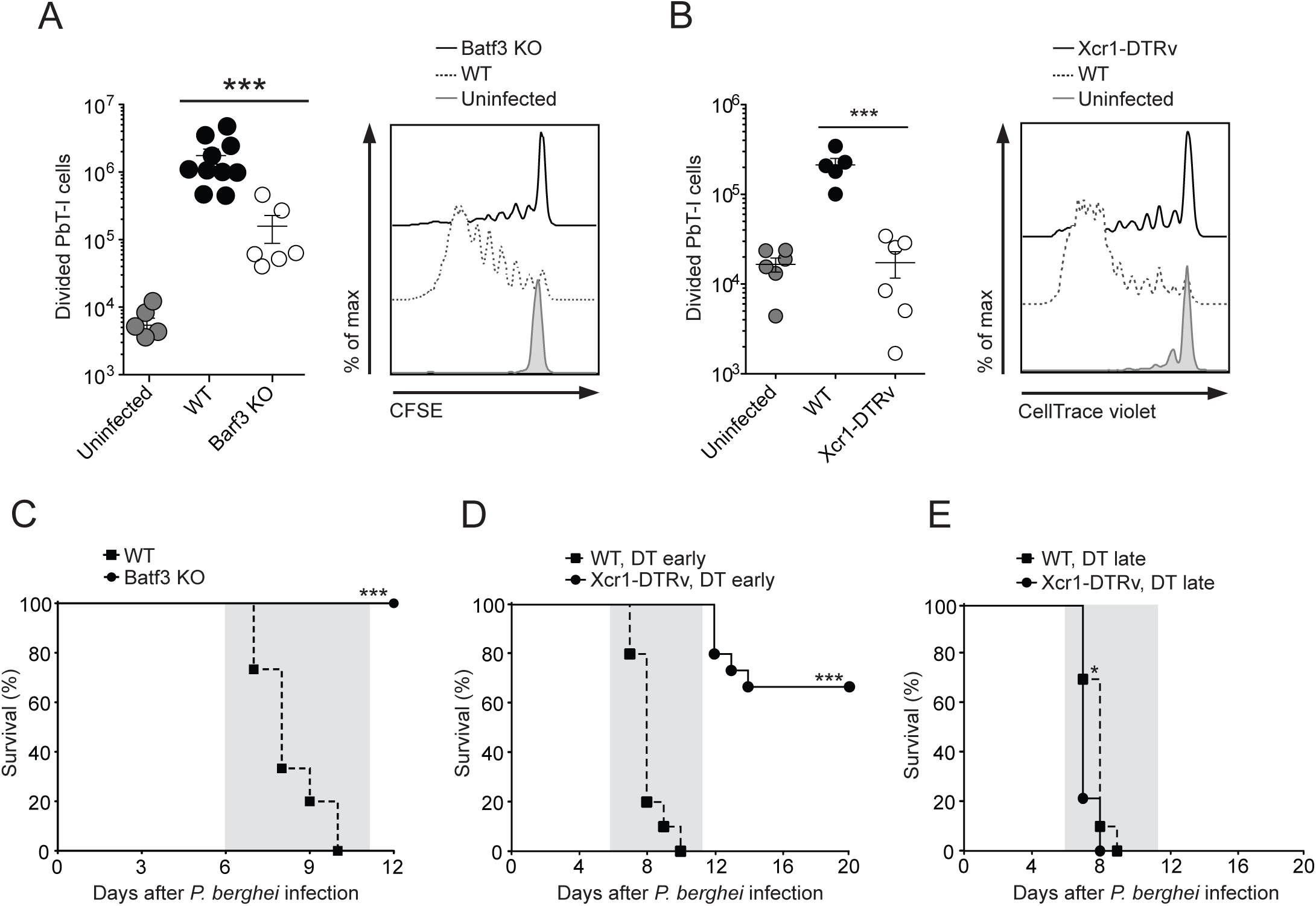
CD8^+^ DC are critical for CD8+ T cell priming and ECM development after PbA infection. (A) PbT-I proliferation in Batf3-deficient mice. WT B6 or Batf3-deficient (Batf3 KO) mice were infected with 10^4^ PbA iRBC and 2 days later 10^6^ CFSE-labelled PbT-I cells were adoptively transferred. 3 days later, spleens were harvested and analysed for proliferation of PbT-I cells. *Left.* The proliferation of PbT-I cells pooled from 3 independent experiments with 2-3 mice in each experiment. Data were log transformed and PbT-I proliferation in infected WT vs Batf3-deficient groups was compared using an unpaired t test (^***^, P<0.001). WT mice (infected and uninfected) include mice homozygous or heterozygous for functional Batf3. *Right.* Representative histogram overlay showing the proliferation of PbT-I cells in WT B6 and Batf3-deficient mice. (B) PbT-I proliferation in DT-treated PbA-infected XCR1-DTRvenus mice. 10^6^ CTV-labelled PbT-I/uGFP cells were transferred into WT or XCR1-DTRvenus mice one day before infection with 10^4^ PbA iRBC. Mice were treated with 500 μg DT i.p. on days -1, 0, 1 and 2 and were killed on day 5 after infection. *Left.* Numbers of divided PbT-I cells in the spleen. Data were pooled from 2 independent experiments. For statistical analysis, data were log transformed and PbT-I proliferation in infected WT vs XCR1-DTRvenus groups was compared using an unpaired t test (^***^, P<0.001). *Right.* Representative histograms showing PbT-I cell division. In A and B, each data point represents the number of proliferating PbT-I cells per spleen and the error bars represent SEM. (C) ECM development in Batf3-deficient mice. WT B6 (dotted line) or Batf3-deficient (solid line) mice were infected with 10^4^ PbA-iRBC and the development of ECM monitored. The shaded area on the graph represents the period in which mice developed ECM. The difference in the survival between the WT and the Batf3-deficient mice is statistically significant (^***^, P<0.001) as determined by the Log-Rank test. Data were pooled from 2 independent experiments including 15 WT and 7 Batf3-deficient mice. (D) ECM development in XCR1-DTRvenus mice treated with DT during the early stages of the infection. Onset of ECM was monitored in WT (dotted line) and XCR1-DTRvenus (solid line) mice infected with 10^4^ PbA-iRBC and treated with 500μg DT on days -1, 0, 1 and 2 p.i. Curves were statistically compared using a Log-rank test (^***^, P<0.0001). Data were pooled from 2 independent experiments including 10 WT and 15 XCRl-DTRvenus mice. (E) ECM development in PbA-infected Xcr1-DTRvenus mice treated with DT in a late stage of the infection. The development of ECM was monitored in WT (dotted line) and XCR1-DTRvenus (solid line) mice infected with 10^4^ PbA-iRBC and treated with 500μg DT on days 5, 6, 7 and 8 p.i. Curves were statistically compared using a Log-rank test (^*^, P=0.0159). Data were pooled from 2 independent experiments including 10 WT and 14 XCR1-DTRvenus mice.

To assess the role of CD8^+^ DC in ECM, we first assessed ECM in Batf3-deficient mice after infection with 10^4^ PbA iRBC (Fig 7C). In agreement with previous reports (67), Batf3-deficient mice did not develop ECM, demonstrating an essential role for CD8^+^ DC in promoting pathology. To obtain a precise indication of the temporal requirement for CD8^+^ DC, we treated XCR1-DTRvenus mice with DT during the early stages of PbA infection (days -1 to 2pi) and monitored for ECM. While non-depleted mice rapidly succumbed to ECM, DT-treated mice were largely protected (Fig 7D), despite harbouring similar levels of parasitaemia (Fig S4A). To determine whether CD8+ DC function may be required during the effector phase of ECM, XCR1-DTRvenus mice were infected with PbA and treated with DT from day 5 post-infection (Fig 7E). In contrast to mice depleted of DC from the beginning of infection, all mice depleted just prior to ECM onset showed rapid disease onset supporting the view that CD8^+^ DC were essential for the priming but not the effector phase of ECM.

### CD4+ T cell help acting on CD8+ DC is required for CD8+ T cell priming during blood stage PbA infection

Primary CD8^+^ T cell responses can be CD4^+^ T cell dependent (18) or independent (19-21). To study the relevance of CD4^+^ T cell help during blood stage malaria, we looked at the expansion of CD8^+^ PbT-I cells in the absence of CD4^+^ T cells. We transferred 1 million CTV-labelled PbT-I cells into MHC II-deficient mice, devoid of CD4+ T cells, and infected them with 10^4^ PbA iRBC. PbT-I proliferation 5 days later was significantly reduced in MHC II-deficient mice compared with their WT counterparts (Fig 8A). Because elevated numbers of CD8^+^ T cell precursors can reduce helper requirements (74), transfer of a high number of PbT-I cells (1 million per mouse) in the previous experiment might have resulted in an artificial enhancement of T cell proliferation in the mice lacking CD4 T cells. To clarify this point, we transferred lower numbers of PbT-I cells (10^5^, 10^4^) into MHC II-deficient mice, infected them with 10^4^ PbA iRBC and examined proliferation on day 5. This revealed that for this infection model, the response of PbT-I cells was similarly reduced in MHC II-deficient mice compared with WT counterparts regardless of the initial number of PbT-I cells transferred (Fig 8B).

**Fig 8.**
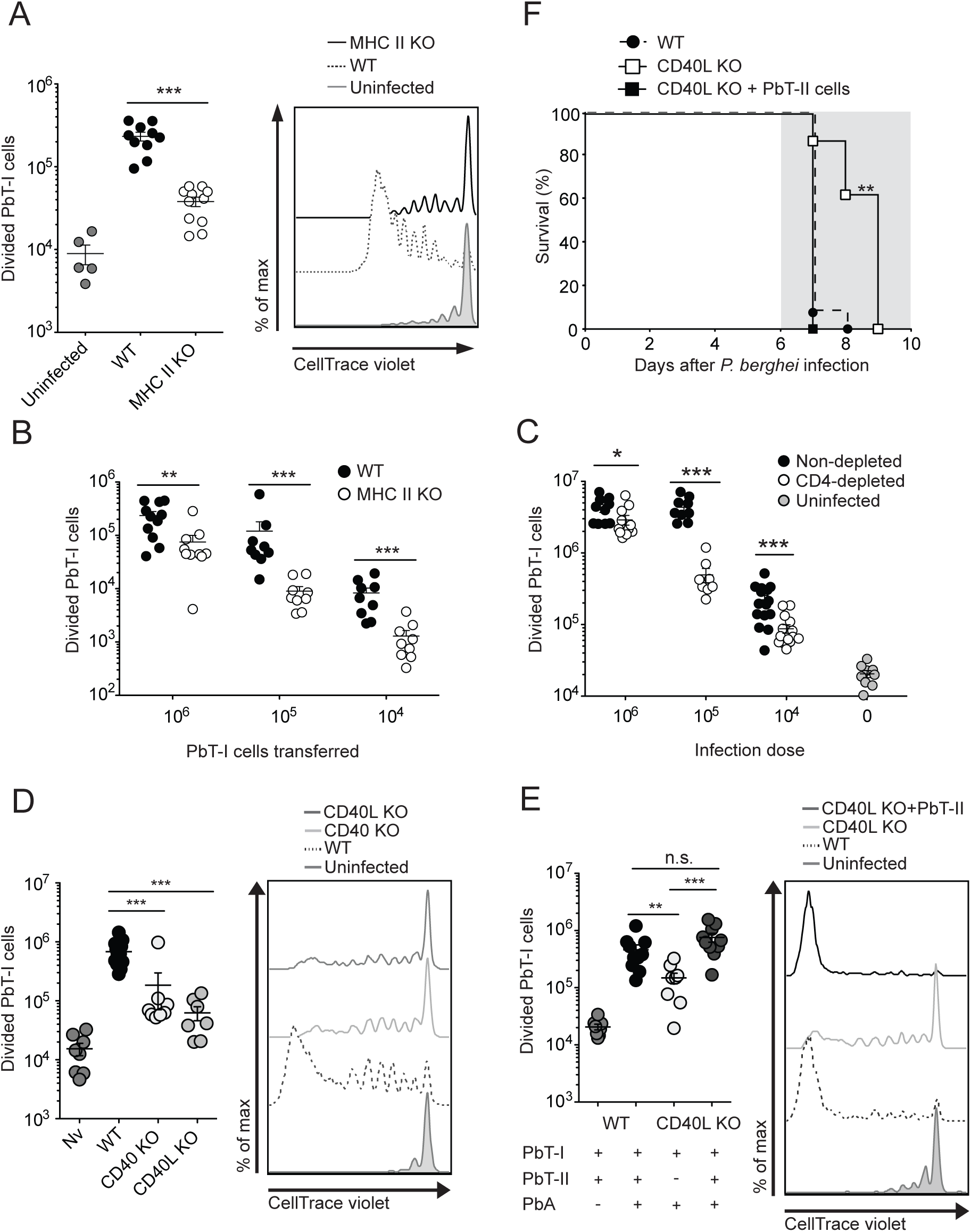
PbT-I priming during blood stage PbA is enhanced by CD4^+^ T cell help via CD40-CD40L interactions. (A) CD4^+^ T cell help is required for PbT-I proliferation. 10^6^ CTV-labelled PbT-I/uGFP cells were transferred into WT or MHC II-deficient (MHC II KO) mice one day before infection with 10^4^ PbA iRBC. Numbers of divided PbT-I cells were assessed on day 5 after infection. *Left.* Numbers of divided PbT-I cells in the spleen. *Right.* Representative histograms showing PbT-I cell division. (B) CD4^+^ T cell help is required independently of the starting number of PbT-I cells. 10^6^, 10^5^ or 10^4^ CTV-labelled PbT-I/uGFP cells were transferred into WT (black circles) or MHC II-deficient (white circles) mice, which were infected with 10^4^ PbA iRBC one day later. Numbers of divided PbT-I cells in the spleens were assessed on day 5 after infection. (C) B6 mice were depleted of CD4 T cells by the injection of 100 μg GK1.5 antibody on days -7 and -4 after infection or left untreated and received 10^6^ PbT-I cells one day before infection with 10^6^, 10^5^ or 10^4^ PbA iRBC. Numbers of divided PbT-I cells were estimated on day 5 after infection. Grey: PbT-I numbers in uninfected WT mice. Data were log transformed and analysed using an unpaired t-test (^***^, P<0.001; ^**^, P<0.01; ^*^,P<0.05; ns, not significant). Data were pooled from 2-4 independent experiments. (D) PbT-I proliferation after PbA infection is decreased in CD40L-deficient (CD40L KO) and CD40-deficient (CD40 KO) mice. WT, CD40- deficient or CD40L-deficient mice received 10^6^ CTV-labelled PbT-I/uGFP cells and were infected with 10^4^ PbA iRBC one day later. Numbers of divided PbT-I cells were assessed on day 5 after infection. *Left.* Numbers of divided PbT-I cells in the spleen. *Right.* Representative histograms showing PbT-I cell division. (E) PbT-II cells restored PbT-I proliferation in PbA-infected CD40L-deficient mice. WT or CD40L-deficient mice were adoptively transferred with 10^6^ CTV-labelled PbT-I cells and some mice also received 10^6^ CTV-labelled PbT-II cells. One day later, mice were infected with 10^4^ PbA iRBC. Numbers of divided PbT-I cells were assessed on day 5 after infection. *Left.* Numbers of divided PbT-I cells in the spleen. *Right.* Representative histograms showing PbT-I cell division. Data were log transformed and analysed using One way ANOVA and Tukey’s multiple comparisons test (^***^, P<0.001; ^**^, P<0.01; ^*^,P<0.05; ns, not significant). Data were pooled from 2-3 independent experiments. (F) PbT-II cells accelerate ECM development in CD40L-deficient mice. CD40L-deficient mice received 10^6^ PbT-II cells (black squares, solid line) or no cells (white squares, solid line) and, together with WT control mice (black circles, dotted line), were infected with 10^4^ PbA iRBC one day later. Mice were then monitored for development of ECM. Survival curves of CD40L-deficient mice with or without PbT-II cells were statistically compared using a Log-rank test (^**^, P=0.0011). Data were pooled from 2 independent experiments including 13 WT, 7 CD40L-deficient mice that received PbT-II cells and 8 CD40L-deficient mice that did not receive PbT-II cells.

Since signals derived from parasite material can promote DC licensing (22), we reasoned that infection with higher numbers of parasites might result in a decreased need for CD4^+^ T cell help. To examine this issue, mice depleted of CD4^+^ T cells with the monoclonal antibody GK1.5 (as an alternative to MHC II-deficient mice), were adoptively transferred with 10^6^ CTV-labelled PbT-I cells and then infected with 10^4^, 10^5^ or 10^6^ PbA iRBC. While numbers of divided PbT-I cells were significantly reduced in CD4^+^ T cell depleted mice compared with non-depleted controls for all parasite doses, the proportional difference was largest for the lower doses (Fig 8C). This indicated that the requirement for help during blood stage infection with PbA was somewhat affected by the infection dose.

Licensing of DC for CTL immunity requires the interaction of co-stimulatory molecules between activated CD4^+^ T cells and DC. CD40L (on the CD4^+^ T cell) and CD40 (on the DC) have been shown to mediate DC licensing in several infectious and non-infectious settings (75-77). To determine whether this pathway was relevant for the provision of help by CD4^+^ T cells during blood stage PbA infection, we transferred PbT-I cells into either mice lacking CD40 or CD40L and infected them with 10^4^ iRBC. PbT-I proliferation and total numbers were significantly reduced in the spleens of both CD40-deficient and CD40L-deficient mice relative to WT controls (Fig 8D). Taken together, the results demonstrated a need for CD4^+^ T cells and CD40-CD40L interactions for the development of optimal CD8+ T cell responses during infection with blood stage malaria.

To demonstrate that CD4^+^ T cells provide help via CD40-CD40L interactions to promote optimal CD8^+^ T cell activation during blood stage PbA infection, we adoptively transferred 10^6^ WT PbT-II cells together with 10^6^ PbT-I cells into CD40L-deficient mice (in which endogenous CD4^+^ T cells were unable to license DC) and then infected these mice with 10^4^ iRBC. While PbT-I proliferation was reduced in CD40L-deficient mice, their response was recovered to WT levels in mice that received PbT-II cells (Fig 8E). To assess the capacity of PbT-II cells to help CD8^+^ T cells cause ECM, we examined the influence of PbT-II cells on the onset of ECM in CD40L-deficient mice. While CD40L-deficient mice showed a significant delay in ECM onset compared to WT mice, this delay was alleviated by transfer of PbT-II cells (Fig 8F). Changes in the time frame for ECM development were independent of parasitaemia, which was not significantly altered by the adoptive transfer of PbT-II cells (Fig S4B). Taken together, these results showed that PbT-II cells were able to help CD8^+^ T cells proliferate and cause ECM in CD40L-deficient mice.

In summary, our results indicate that CD8^+^ DC are the main APC priming both CD4^+^ and CD8^+^ T cells during infection with blood stage PbA parasites. Using our newly generated PbT-II transgenic cell line, we demonstrated that optimal expansion of CD8^+^ T cells required CD4^+^ T cell help, which was provided to the DC via CD40-CD40L interactions.

## Discussion

While TCR transgenic mouse lines are very useful for deciphering the mechanisms by which T cells operate and carry out their functions in different contexts of infection, the available range of these lines specific for blood stage malaria antigens is limited. Here we introduce a new CD4+ T cell transgenic mouse line, termed PbT-II, specific for a broad range of *Plasmodium* species and responsive to both blood stage and pre-erythrocytic stage infections, offering wide applicability for the study of CD4^+^ T cell function in multiple malaria models. To date, the only available CD4^+^ T cell receptor transgenic murine line specific for malaria parasites (*P. chabaudi*) was B5, which is restricted for I-E^d^ and therefore only usable in the BALB/c background. The PbT-II mice provide the opportunity to analyse *Plasmodium-specific* CD4^+^ T cells on the B6 background, for which availability of reagents and gene deficient mice is much broader. We show here that PbT-II cells are an excellent tool for studying CD4^+^ T cell function in the context of malaria immunity or pathology. When adoptively transferred into CD40L-deficient mice, in which endogenous CD4^+^ T cells are unable to fully activate B cells or DC (48, 75, 77), PbT-II cells were able to replace endogenous CD4^+^ T cells for the generation of both antibody responses, which allowed control of *P. chabaudi* parasitaemia, and CD8^+^ T cell responses, which caused ECM after PbA infection. This indicates that PbT-II cells are an invaluable tool for the study of CD4^+^ T cell biology during infection with *Plasmodium* parasites.

The specific antigen recognised by PbT-II cells is yet to be defined. The broad crossreactivity to different *Plasmodium* species suggests that the target epitope belongs to a conserved protein, likely to have an essential function in blood stage parasites. The PbT-II target protein is also expressed during the pre-erythrocytic stage, as PbT-II cells also responded to RAS. It is possible that this protein is only expressed late during the liver stage, shortly before merozoites are released to the blood (78), allowing for a small window of antigen presentation during the liver stage of the parasite. However, the limited development of RAS within hepatocytes argues against this possibility. Alternatively, the abundance of this protein during the liver stage of the parasite cycle may be lower than during the blood stage. Further studies will be required to determine the exact antigen recognized by PbT-II cells.

DC are extremely efficient at initiating T cell responses in numerous infection models. We have been able to demonstrate a pivotal role for the CD8+ DC subset in the priming of both CD4^+^ and CD8^+^ T cells during blood stage PbA infection. Our results on CD8^+^ T cell activation confirm previous *ex vivo* data showing a superior efficiency of CD8^+^ DC at priming OT-I cells after exposure to transgenic PbA parasites expressing SIINFEKL (9). They are also consistent with *in vivo* studies showing a reduced number of activated endogenous CD8^+^ T cells after PbA infection of mice deficient in CD8+ DC (37, 38, 67). As CD8^+^ DC possess specialised machinery for cross-presentation (34), this likely underlies their unique capacity to prime CD8+ T cells during infection by *Plasmodium* species.

CD8^+^ DC were also the main subset that stimulated D78 and PbT-II cells. To estimate the net contribution of the different DC subsets to MHC II presentation, the results obtained for D78 stimulation *ex vivo* (Fig 5B), which show that CD8^+^ DC are about 10-fold better than CD4^+^ DC or DN DC at presentation, were adjusted to reflect the distinct abundance of sorted DC subsets in the spleen (CD4^+^ DC, 55.1%; CD8+ DC, 17.7%; DN DC, 13.4% of splenic CD11c^hi^ MHC II^hi^ cells). From this adjustment, we estimated that the contribution of CD8+ DC to antigen presentation via MHC II was 67% of the total, whereas that of CD4^+^ DC contributed 20% and DN DC contributed the remaining 13% (with the combined CD8^−^ DC subsets therefore contributing 33% of the total antigen presentation in the spleen). *In vivo* experiments using different mouse lines deficient in CD8+ DC to assess the contribution of CD8^+^ DC to PbT-II priming during blood stage PbA infection supported our observations with the D78 hybridoma: the number of dividing PbT-II cells in the spleen of Batf3-deficient or IRF8-deficient or DT-treated XCR1-DTRvenus mice was reduced to 45.1%, 16.6% and 31.0%, respectively. Because of the limitations of Batf3-deficient and IRF8-deficient mice in this system, with Batf3-deficient mice still containing immature DC capable of MHC II presentation (68, 69), and IRF8-deficient mice presenting additional immune defects (44, 70), the outcome using XCR1-DTRvenus mice likely best represents the net contribution of CD8^+^ DC to MHC II presentation. This value (69%) very closely aligns with the 67% contribution to MHC II presentation estimated using the D78 hybridoma.

These results clearly define CD8^+^ DC as the major APC contributing to CD4+ T cell priming during PbA infection. Previous work in our lab showed roughly equivalent CD4^+^ T cell priming *ex vivo* by CD8^−^ and CD8^+^ DC from BALB/c mice infected with a transgenic PbA parasite expressing various model T cell epitopes (9). In that case, responses in B6 mice could not be assessed because they were too weak (9). Consistent with our current study, other reports have found an important role for CD8^+^ DC in CD4^+^ T cell activation: Clec9A-DTR mice depleted of CD8^+^ DC by injection of DT showed a markedly decreased activation of endogenous CD4+ T cells during PbA infection (37). Also, CD8+ DC were more efficient than CD8^−^ DC at presenting *P. chabaudi-derived* antigens to CD4^+^ T cell hybridomas (39). In the latter study, the superiority of CD8^+^ DC was only observed for DC derived from uninfected mice, and CD8^−^ DC extracted on day 7 after infection were more efficient at MHC II presentation than CD8^+^ DC. At that late time point, CD8^+^ DC showed greater cell death than CD8^-^ DC, which likely explained their decreased stimulatory capacity (39).

We also provided evidence that CD8^+^ DC contribute to the quantity of the T cell response, as CD8^+^ DC depletion led to the generation of reduced frequencies of Th1 and Tfh cells. This agrees with existing literature showing that the CD8^+^ DC subset preferentially generates Th1 responses over Th2 responses (35, 36) and is efficient at promoting Tfh immunity (71, 72). The consequences of this functional capacity for CD8^+^ DC, however, are likely to differ depending on the model studied. During PbA infection, for example, IFN-γ producing CD4^+^ T cells have been implicated in ECM by attracting CD8^+^ T cells to the brain (79). Thus, loss of Th1 cell development after CD8^+^ DC depletion may impact CD8^+^ T cell recruitment to this organ; however, in this example CD8^+^ DC would also be crucial for initiation of CD8^+^ T cell responses. Alternatively, control of *P. chabaudi* infection requires Th1 cells during the first peak of parasitemia (80) and Tfh cells for complete elimination of the parasite (81). Not surprisingly, the absence of CD8+ DC in this infection results in impaired parasite control and more pronounced relapses (40).

The reason for the observed dominance of CD8^+^ DC in MHC II presentation remains unclear. The form of antigen encountered by DC appears to play a role, as in the present work CD8^+^ DC only out-performed other DC subsets when antigen was provided in a RBC- associated form, but not when in soluble form (Figs 5C and D). This is consistent with their specialization in the capture of cell-associated antigen (82). Further studies are required to elucidate the mechanisms underlying the superiority of CD8^+^ DC in MHC II presentation during PbA infection.

Given the critical role of CD8^+^ T cells in the development of ECM (83), determining the requirements for CD8^+^ T cell activation during blood stage malaria is of great importance. Conventional DC are essential for the induction of ECM (66) and lack of ECM in XCR1- DTRvenus mice treated with DT early in infection confirms the findings of others (37) that implicate CD8^+^ DC in this process. This is not surprising given their central role in priming both CD4^+^ and CD8+ T cells. Less clear is the extent at which CD4^+^ T cell licensing of CD8^+^ DC is required for ECM development. Several studies have shown that anti-CD4^+^ antibody given around the time of PbA infection prevents ECM (26-30), consistent with the view that CD4^+^ T cell help is required for the induction of CTL responses. However, CD4^+^ T cells have also been implicated in CD8^+^ T cell recruitment to the brain, thus raising the possibility that their depletion merely affects this facet of disease. Only one study provides direct evidence for a contribution of CD4^+^ T cell help to CD8^+^ T cell proliferation, where CD4+ T cell depletion resulted in low proliferation of OVA-specific OT-I cells after infection with transgenic PbA parasites expressing the OVA epitope (31). By using our PbT-I transgenic CD8^+^ T cells, we clearly demonstrated that CD4+ T cell help was essential for optimal CD8^+^ T cell expansion, as mice depleted of CD4^+^ T cells showed suboptimal PbT-I proliferation. These responses were dependent on CD8^+^ DC, as they were impaired when this subset of DC were depleted. Furthermore, they depended heavily upon CD40L, as mice deficient in this molecule had curtailed CD8^+^ T cell proliferation. The capacity of transferred PbT-II cells to help PbT-I cells to proliferate in CD40L-deficient mice further implicated CD40L as an important component of help that could be provided by PbT-II T cells. Interestingly, CD40 signalling was not absolutely essential for ECM development, as CD40L-deficient mice eventually developed ECM in the absence of added PbT-II T cells, though with a delayed kinetics. This contrasts a previous report that found CD40L- deficient mice were resistant to ECM (84), an observation that may be explained by differences in animal housing facilities, potentially microbiota, known to affect ECM (85).

Our work extends earlier findings by determining that CD4^+^ T cell help is required for optimal CD8^+^ T cell expansion after infection with different doses of parasites, and is relatively independent of the initial CD8+ T cell precursor frequency. By defining CD40 signalling as an important event in DC licensing in this model, we have also identified a key checkpoint that could be targeted to limit pathogenic CD8^+^ T cell immunity to blood stage disease.

We have established that CD8^+^ DC are the main APC for CD4^+^ and CD8^+^ T cell priming, and that CD8^+^ T cell priming required CD4^+^ T cell help. It is likely that CD8^+^ DC are licensed by those CD4^+^ T cells that they prime, though CD4^+^ T cells primed on CD8^-^ DC may also contribute to this process. The essential role for CD8^+^ DC in CD8^+^ T cell priming and their dominant role in CD4^+^ T cell priming puts this subset at the centre of response initiation to blood stage malaria parasites and suggests that strategies aimed at generating T cell responses against blood stage malaria parasites will benefit from exploiting this DC subtype. The findings that individuals with severe disease in malaria endemic areas present with increased numbers of circulating BDCA3^+^ DC (86) (the human equivalent to mouse CD8^+^ DC) and activated CD8^+^ T cells (38) relative to those with mild malaria, suggest that CD8^+^ T cell activation during blood stage malaria may be driven by a similar process in humans as in the mouse model. More precise knowledge of the mechanisms that drive CD8^+^ DC function during malarial infections may, thus, allow us to develop more effective strategies to combat this disease.

